# Data-driven forecasts of regional arrivals of non-native vertebrates worldwide

**DOI:** 10.64898/2026.07.08.737252

**Authors:** César Capinha, Matilde Mendes, Joana Catarino, Filipa Coutinho Soares, Franz Essl, Hanno Seebens, Sandra Oliveira, Luís Reino, Joana Ribeiro

## Abstract

**Aim:** To forecast near-future arrivals of non-native terrestrial and freshwater vertebrates at the regional level.

**Location:** Global (geopolitical regions worldwide, including countries and main administrative divisions).

**Methods:** We compiled first regional record data and assembled functional and macroecological variables for 1,931 non-native vertebrate species. For each region, we identified ‘recently arrived’ non-native species using retrospective windows of thirty and twenty years ending in 2015 (1986–2015; 1996–2015). We then fitted region-specific random-forest models classifying recently arrived species versus those not yet arrived using as predictors: (i) harmonised species traits (e.g., habitat, diet, body size and native-range attributes) and (ii) spread history, capturing time since first record elsewhere. Predictive performance was evaluated using leave-one-out cross-validation, comparing full models with trait-only and spread-only variants. We also assessed relationships between predictive accuracy, predictor importance, and the geographic positioning and trade connectedness of regions. Finally, we predicted region-specific probabilities of arrival for species not yet recorded.

**Results:** Forecasting accuracy was consistently high across regions and taxa, with AUC values above 0.9 in more than half of the focal regions. Full models substantially outperformed models using either predictor set alone, and spread-history-only models typically exceeded trait-only models. Relative importance of spread-history predictors declined with geographic distance to the focal region, whereas predictability was lower in highly trade-connected regions. Predicted near-future high-risk arrivals were dominated by birds and freshwater fishes and showed strong regional structuring. A small set of species ranked highly across many regions (e.g., birds: *Phasianus colchicus, Acridotheres tristis, Amandava amandava, Colinus virginianus, Corvus splendens* and *Lonchura malacca*; fishes: *Coregonus peled* and *Oreochromis mossambicus*; mammal*: Oryctolagus cuniculus*), suggesting substantial unrealised spread potential.

**Main conclusions:** Near-future regional arrivals of non-native vertebrates are predictable from spread history and species traits. This enables scalable, updateable regional watchlists to support prevention, early detection and horizon scanning.

## Introduction

In recent decades, intensified global transport, trade, and human mobility have increased both the frequency and diversity of organisms introduced beyond their native ranges (IPBES, 2023). Although most introductions fail to establish self-sustaining populations, the number of established non-native species continues to increase (Seebens et al., 2025), reshaping regional species pools and global biogeographic patterns (Olden, 2006; Capinha et al., 2015). While only a subset of established non-native species becomes invasive, their impacts can be disproportionate, driving biodiversity loss and ecosystem change, imposing high economic costs, and affecting human health (Pyšek et al., 2020a; Carneiro et al., 2025). The IPBES assessment reports >37,000 established non-native species globally, with ∼200 newly recorded each year, and annual costs exceeding US$423 billion in 2019 (IPBES, 2023). Together, these trends highlight a key need: anticipating where ongoing invasions are most likely to spread in the near future to strengthen prevention, surveillance, and rapid response.

Biological invasions unfold as a strongly structured, interregional process. Analyses of first-record sequences suggest that a small set of highly connected countries tends to record non-native species early, after which spread often proceeds to geographically close-by regions, consistent with a two-level architecture comprising global “backbone” hubs followed by regional radiative spread (Capinha et al., 2023). This structure is further reinforced by secondary introductions and bridgehead dynamics, whereby established non-native populations become additional sources that accelerate spread and generate repeated introductions to other regions (Lombaert et al., 2010; Bertelsmeier et al., 2018; Montgomery et al., 2023). Moreover, spread trajectories can retain a historical signal: past invasion intensity and long-standing connectivity may continue to shape contemporary spread, even where policies and trade patterns have changed (Lenzner et al., 2022). However, regional connectivity alone cannot explain non-native species spread, as pathways (e.g., the pet trade, aquaculture, or game-species releases) are also filtered by species traits (Street et al., 2023). Species traits can influence exposure to human-mediated transport, and thus propagule pressure (i.e., the number and frequency of individuals introduced; Lockwood et al., 2005), as well as the likelihood of overcoming demographic, dispersal, and environmental barriers following arrival (Blackburn et al., 2011; Allen et al., 2017; Gippet and Bertelsmeier, 2021). Together, these patterns imply that invasion histories, especially when combined with trait information, can provide a basis for anticipating where non-native species are most likely to arrive next.

A range of approaches can support forward-looking prioritisation, but each faces constraints when the goal is near-future spread forecasting across many species and many regions. Species distribution models and related niche-based approaches can identify areas of potential environmental suitability, but forecasting ongoing invasions is challenging under non-equilibrium dynamics, dispersal limitation, changing networks of trade and environmental change, and shifting realised niches (Gallien et al., 2012; Pili et al., 2020). Pathway- and network-based models can represent human-mediated transport more directly and have produced robust forecasts where movement data are well characterised (e.g., Montgomery et al., 2023; Oliveira et al., 2023; Zhang et al., 2024). Expert-led horizon scanning is likewise widely used to prioritise high-risk species expected to arrive and establish within a defined time horizon (e.g., Kenis et al., 2022; Oficialdegui et al., 2023). For example, a revisit of a Great Britain horizon scan found that 31 of 92 shortlisted species had arrived by 2023, including 12 of the top 20 within 10 years (Peyton et al., 2026). Nonetheless, scale remains a persistent limitation, with many predictive studies and assessments designed for single taxa, individual pathways, or a limited set of regions, even as the candidate pool of potential invaders continues to grow (Seebens et al., 2018). In addition, the expertise and resources required for repeated horizon scans and sustained surveillance are unevenly distributed, leaving many high-exposure regions with limited capacity for proactive prioritisation.

Here we present a complementary, data-driven approach designed explicitly for broad geographical and taxonomic coverage: region-by-region estimates of the probability of near-future “arrival” across many species, using information embedded in observed invasion histories and species traits. Building on global first-record data for terrestrial non-native vertebrates, we fit region-specific prediction models in which recent first records (within a retrospective calibration window) are treated as evidence of arrival, and explained by two information streams: (i) harmonised functional and macroecological species traits linked to transport exposure and post-arrival success; and (ii) explicit measures of prior spread elsewhere (i.e., time since first record in other regions), reflecting the spread structures documented at global scales. The resulting models yield probabilistic, region-level rankings of species according to their likelihood of arrival, enabling updateable regional watchlists that complement, rather than replace, pathway analyses and expert-led horizon scanning. Finally, we evaluate predictive performance and examine how the importance of spread-history information varies with geography, helping to clarify when and where invasion history is most informative for near-term forecasting.

## Data and Methods

### Identification of focal species and first record data

To develop our models (which rely on non-native spread-history data; see below), we required information on the timing of the first regional record of non-native species. For this purpose, we used the FirstRecords Database v3.1 (Seebens, 2023), which provides the earliest known year of recording of non-native species in the wild across regions worldwide (countries and subnational units such as provinces or islands; hereafter simply ‘regions’). It is the most up-to-date and comprehensive resource of its kind, containing over 75,000 first records for more than 24,000 non-native species across 296 regions. The database includes records of established, non-established, eradicated, and extinct non-native species in both countries and subnational regions. Because we aimed to analyse patterns of species spread, we retained all types of records, irrespective of invasion status.

We focused on terrestrial and freshwater vertebrate groups (amphibians, birds, mammals, reptiles, and fishes). Marine species were excluded because their dispersal can occur largely outside terrestrial political boundaries, making geopolitical regions unsuitable units for modelling spread. To identify and exclude marine fishes, we used FishBase habitat designations (freshwater/brackish/saltwater occurrence fields) (Froese and Pauly, 2025). A small number of marine mammals (e.g., seals, sea otters, and manatees) and seabirds (e.g., penguins, frigatebirds, and albatrosses) were also manually excluded.

### Functional and macroecological trait variables

To use as predictors of non-native species spread, we compiled for each species a set of functional and macroecological trait variables describing: (1) habitat, (2) diet, (3) body size and (4) characteristics of species’ native ranges (Table 1; Figure 1). This trait set aims to capture both the propensity of species to enter human-mediated transport pathways and their ability to overcome the main demographic and environmental barriers along the introduction-establishment-spread continuum (Blackburn et al., 2011). For example, large-bodied, generalist vertebrates with broad native ranges and relatively fast life histories tend to be over-represented among species traded as pets or introduced for hunting (Blackburn et al., 2017; Street et al., 2023). Likewise, species whose native ranges already encompass human-dominated landscapes are more likely to be transported and to find suitable conditions in recipient regions (Carlon and Dominoni, 2024), consistent with macroecological frameworks that link invasion success to the interplay between species traits, source-area characteristics and introduction pathways (Pyšek et al., 2020b).

**Figure 1.**
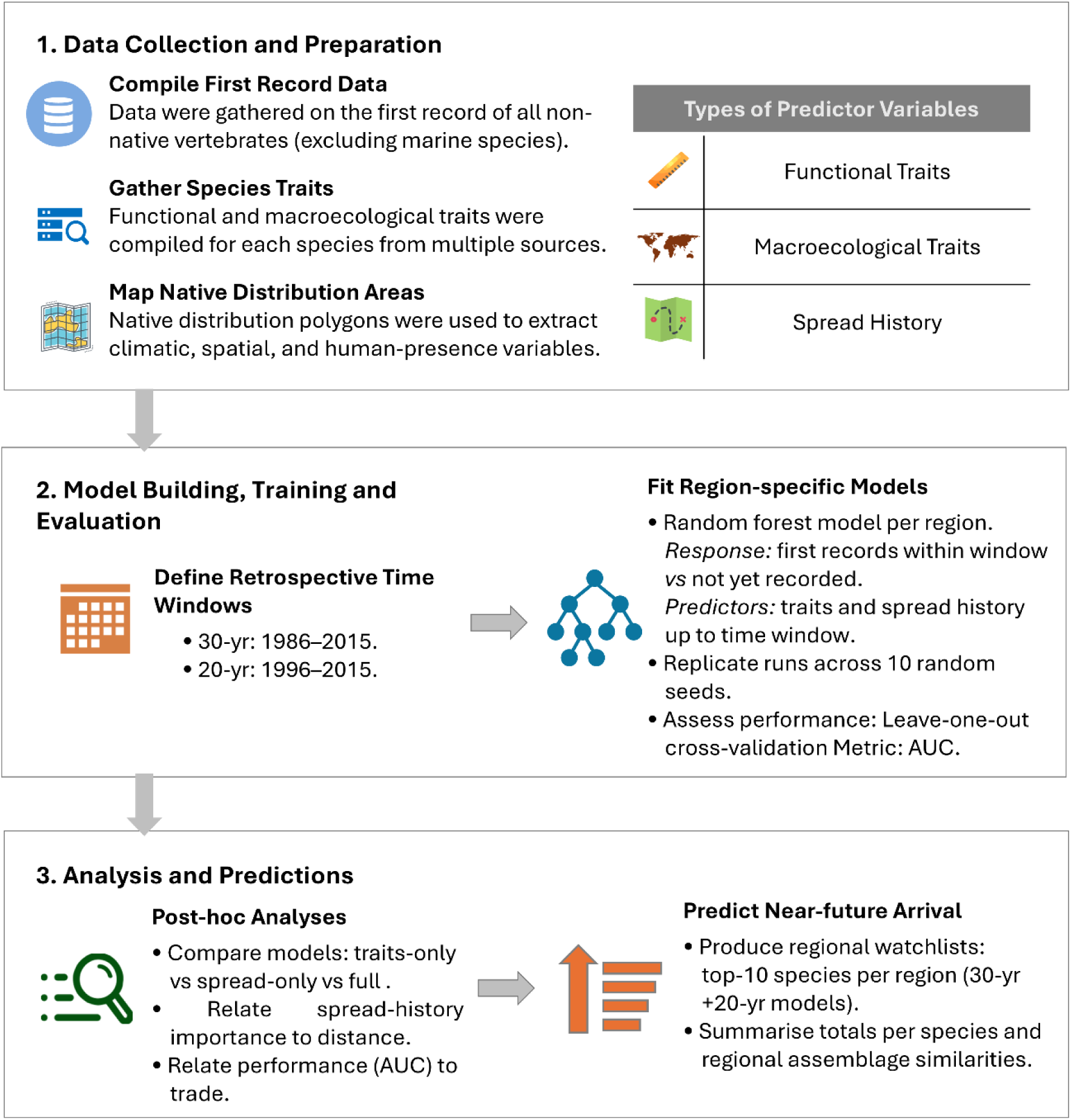
Main steps of the modelling approach. The workflow includes data collation and processing, model development, training and evaluation, and analysis of results. Predicted probabilities are then used to forecast regional arrivals of non-native vertebrates

**Table 1.**
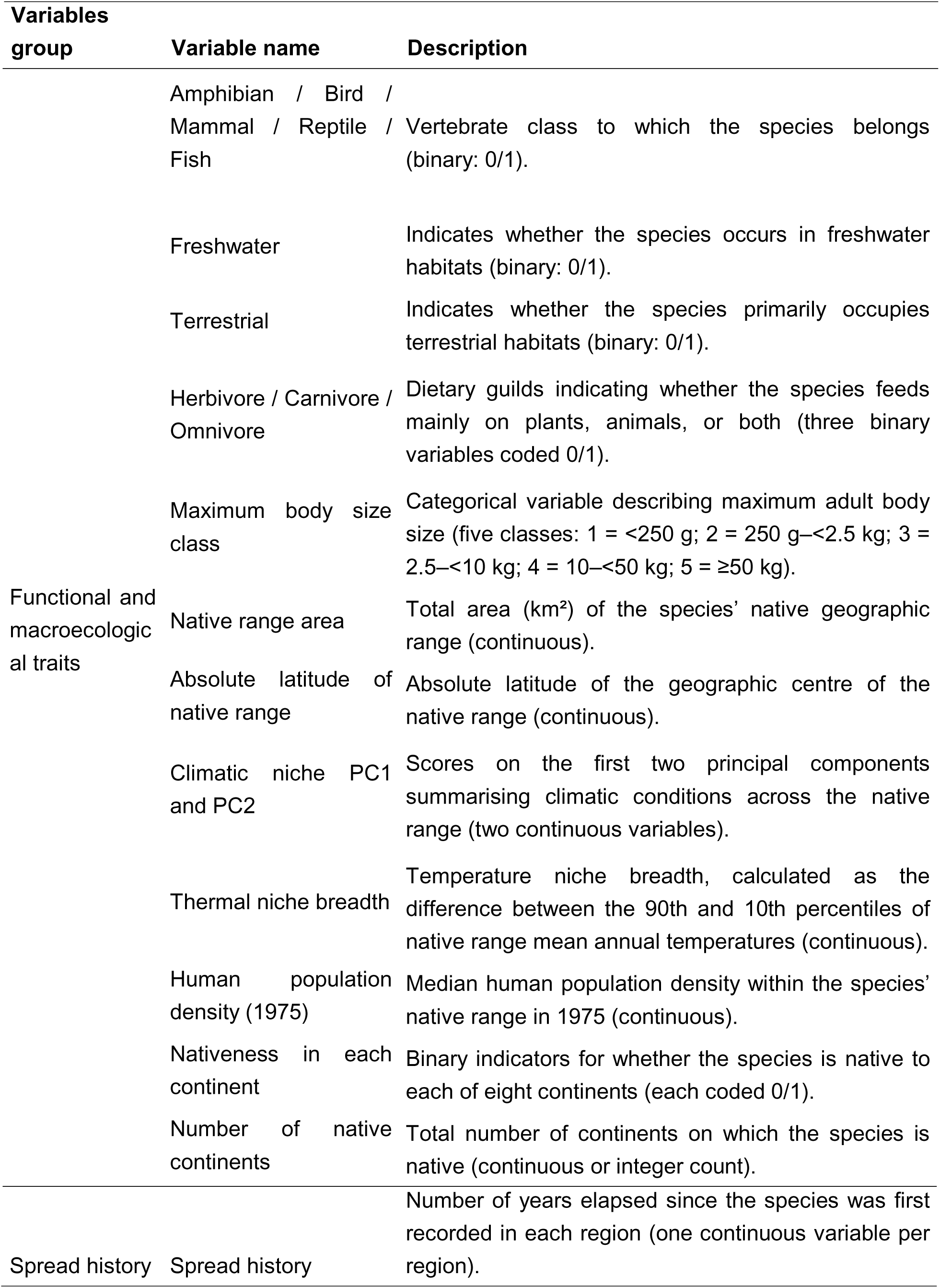
Functional and macroecological traits and spread-history variables used as predictors in the models.

Data on habitat, diet and body size were compiled from taxon-specific trait databases: BirdLife International (Birdlife, 2025) and AVONET (Tobias et al., 2022) for birds; AmphiBIO (Oliveira et al., 2017) for amphibians; PHYLACINE (Faurby et al., 2018) for mammals; ReptTraits v1.2 (Oskyrko et al., 2024) for reptiles; and FishBase (Froese and Pauly, 2025) plus ITOFF (Invasive Traits of Freshwater Fish; Jessop et al., 2023) for fishes. Missing values in these sources were filled, where available, through literature searches.

For amphibians and reptiles, primary trait data sources reported maximum adult body mass, whereas for birds and mammals, they provided mean body mass. To make these measures comparable, we converted mean masses for birds and mammals to maximum masses using empirically derived scaling factors of 1.15 and 1.33, respectively (Dunning, 2007; Prothero, 2015). For fishes, only the maximum body length was comprehensively available. We converted length (L) to mass (weight, W) using the allometric relationship (W = aL^b) with general parameters (a = 0.0105) and (b = 3.0) within empirically reported ranges (Froese, 2006; Simon et al., 2023). To reduce sensitivity to uncertainty in length–mass conversions, we discretized maximum body mass into five broad ordinal size classes (1–5): <250 g, 250 g–<2.5 kg, 2.5–<10 kg, 10– <50 kg, and ≥50 kg.

To derive macroecological traits of species’ native ranges, we compiled spatial data on native ranges of amphibians and mammals from the IUCN Red List (IUCN, 2025), birds from BirdLife (Birdlife, 2025), reptiles from the Global Assessment of Reptile Distributions dataset (Roll et al., 2017), and freshwater fishes from Tedesco et al. (2017). For a small number of mammal species missing from the IUCN polygons, we supplemented with PHYLACINE 1.2 current range polygons and information from the primary literature. For fishes, the best-available data are at the river sub-basin level (Tedesco et al., 2017); we therefore used sub-basins identified as native and with occurrence status classified as “valid” (i.e., excluding questionable occurrences). Across all taxa and sources, we retained only polygons corresponding to native or reintroduced ranges (i.e., former native areas where the species has re-established) and removed all non-native, vagrant, uncertain and assisted-colonisation areas.

From these native-range polygon layers, we calculated a set of spatial predictors in R (R Core Team, 2024). Total native range area (km²) was obtained by projecting each species’ polygons to an equal-area coordinate system (Equal-Area Scalable Earth Grids; EPSG:6933) and summing polygon areas.

We also overlaid native ranges with global climate data WorldClim v2.1, 30 arc-second resolution variables (Fick and Hijmans, 2017). To characterise thermal niche breadth, we performed a principal component analysis, with the first two climatic components explaining 82% of the total variance, based on the 10th, 50th and 90th percentiles of annual mean temperature, annual precipitation, temperature seasonality and precipitation seasonality across each native range.

Degree of human presence within native ranges was quantified by overlaying species’ native range polygons with gridded human population density from the History database of the Global Environment (HYDE) dataset for the year 1975 (Goldewijk et al., 2011), from which we extracted the median population density. In a small number of cases, mainly small islands, HYDE coverage was unavailable. For these regions, we consulted the literature and used the population density estimates from the closest year available.

To characterise native-continental boundaries, we intersected each species’ native range with a categorical raster of the world’s continents (ESRI, 2024). For each species, we recorded presence/absence on each continent (yielding eight binary “nativeness” variables) and calculated the total number of continents on which the species is native.

Finally, to derive the geographic centre of the native range, we rasterised each native range polygon onto a template grid (matching the climate data) and calculated the mean latitude of all occupied cells. The absolute value of this mean latitude was used as a measure of the species’ native latitudinal position.

### Spread history variables

In addition to the functional and macroecological variables, we calculated for each species the number of years elapsed since its first record in each region (i.e., residence time). These values were calculated relative to the start year of the time window used in the models (see below). Species with no prior records in a given region were coded as 0 (i.e., not yet recorded).

Species with missing data for one or more variables were excluded from subsequent modelling analyses. In total, we obtained complete information for all trait and spread history variables for 1,931 species, representing >97% of all non-marine vertebrates in the FirstRecords Database v3.1. The retained dataset comprised data from 276 regions and a total of 8,831 first-record events.

All spatial analyses were performed in R using the packages *sf* (Pebesma and Bivand, 2023), *terra* (Hijmans et al., 2022), *exactextractr* (Baston et al., 2021) and associated functions (R Core Team, 2024).

### Modelling procedure and performance assessment

Our main aim was to estimate the probability of a given non-native vertebrate species arriving in each region in the near future. Here, ‘arrival’ refers to a first record in the focal region, regardless of the pathway (e.g., deliberate or accidental human-mediated introduction, or secondary spread from non-native populations established elsewhere; sensu Peyton et al., 2026). To do so, we implemented a region-specific modelling framework, fitting a separate predictive model for each region (Figure 1). In each model, the response variable indicates whether a species was first recorded within a recent time window (*positives*, coded as 1) or not (*negatives*, coded as 0). To predict this binary outcome, models used variables describing species’ functional and macroecological traits and their prior regional spread history (described above) (Fig. 2). After training, the models were applied to estimate future arrival probabilities for each non-native species under the current spread history.

**Figure 2.**
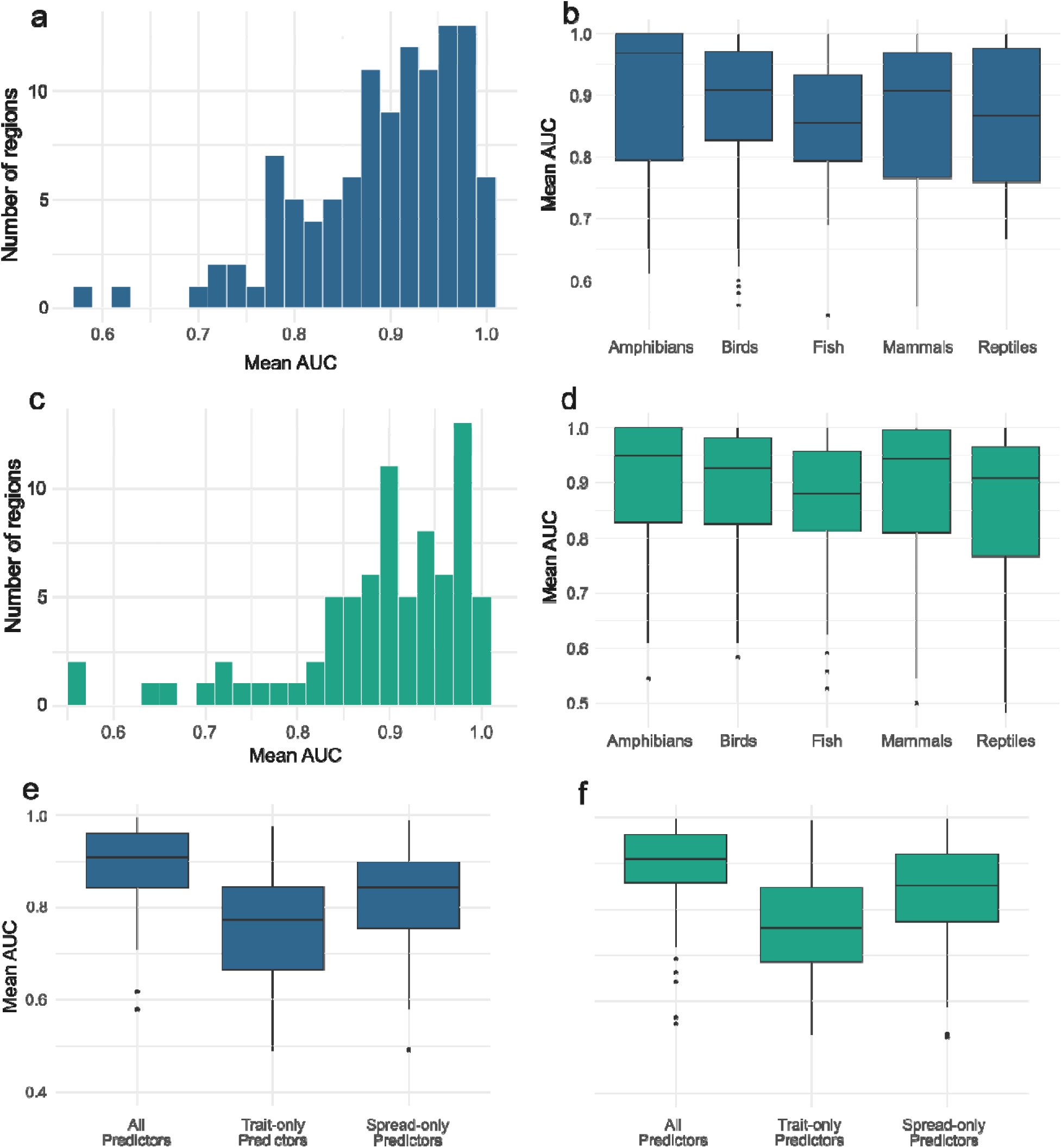
Variation in AUC values across regions, comparing models calibrated with a 30-year period (blue) and a 20-year period (green). Models were built using different predictor subsets: the full set of variables (All), only functional and macroecological traits (Trait-only), or only spread history (Spread-only). For each model, AUC values correspond to the average across ten repetitions to account for the stochasticity of the random forest algorithm.

A key consideration in defining the response variable is the selection of the retrospective time window used to represent ‘recent introductions’. This choice requires balancing two aims: (i) maximising the number of positive records available for each region to support model fitting, and (ii) ensuring that the time window represents a period that is sufficiently recent to reflect current spread dynamics. Because reporting delays may affect the most recent years of the FirstRecords Database (Seebens et al., 2017; Brock and Daehler, 2025), including the latest data year could introduce bias. For example, in our dataset, 554 first records fell within 2005-2015, whereas only eight were reported after 2015, a distribution indicative of substantial reporting lags. Therefore, 2015 was set as the end year of the retrospective window.

Models could only be fitted for regions providing a minimally sufficient number of positive cases (i.e., non-native species recorded within the time window) and negative cases (non-native species not recorded in the region). Negative cases were abundant across all focal regions, whereas positive cases were more limited. Based on preliminary assessments, we defined a minimum of five positive records, for a region to be included in the modelling analyses.

To help define the length of the retrospective windows, we evaluated cumulative 10-year increments (counting backwards from 2015) and quantified, for each window, the number of regions containing five or more positive records. This procedure revealed a clear trade-off between temporal recency and the number of regions that could be modelled (Fig. S1 in Supplementary Material). To balance these considerations, we opted for using two retrospective windows: a 30-year window from 1986 to 2015, which allowed model fitting for 110 regions (hereafter the ‘30-year’ period), and the 20-year window from 1996 to 2015 (‘20-year’ period), which included fewer regions (n = 77) but focused on a more recent period, better aligned with contemporary spread dynamics (Figs S2 and S3).

In each model, to ensure that negative records reflected non-native species that had not yet arrived, but could plausibly do so, we excluded all non-native species with first-record dates earlier than the start of the window (i.e., those that were introduced, but before the time period of reference). We also excluded species native to the region based on SInAS v3.1 dataset (Gómez-Suárez et al., 2025), which provides harmonised global data on species’ native and non-native countries worldwide, as well as non-native species lacking a known first-record year (i.e., introduced but not known when). These filtering steps ensured that the dependent variable for each focal region included only: (i) non-native species first recorded in the time window of interest (positives), and (ii) non-native species not yet recorded (negatives).

As predictors of this variable, we used species’ functional and macroecological traits together with variables capturing their spread history (explained above; Table 1). Prior to model fitting, to avoid collinearity issues, we calculated Spearman correlations among predictors and excluded variables with |ρ| ≥ 0.7 using the *findCorrelation* algorithm in *caret* (Kuhn, 2008).

### Model fitting and evaluation

We then fitted region-specific models for predicting species’ probability of arrival using random forests (Breiman, 2001). The choice for this algorithm was partly motivated by the consistently strong predictive performance of random forests and the ability to deal with large numbers of predictors (Breiman, 2001; Valavi et al., 2022), but also because it offers comparatively fast computation times relative to most other machine learning approaches (Valavi et al., 2022). This latter aspect was of particular significance in the context of this work, which required fitting models across a large number of regions with multiple replicates to quantify algorithmic stochasticity, evaluate model performance, and conduct post-hoc analyses (see below).

The models were fitted using the function *imbalanced* of the package *randomForestSRC* (Ishwaran et al., 2021). This function addresses the issue of class imbalance (i.e., in our case, generally lower numbers of positives than of negatives) by internally reweighting and resampling the minority class. For model fitting, we used 3,000 trees, a number sufficiently large to ensure the representativeness required for resampling procedures. For each predictor variable, we also calculated permutation- based importance, quantified as the increase in out-of-bag prediction error after randomly permuting that predictor’s values relative to the non-permuted model (Ishwaran et al., 2021).

To evaluate the predictive performance of each model, we used leave-one-out cross-validation (LOOCV). LOOCV was implemented by iteratively holding out one species from the evaluation set, refitting the model without the withheld species, and predicting its probability of arrival. Given the imbalance between classes (negatives outnumbering positives), we built a balanced evaluation set by testing all positive records and randomly sampling the same number of negatives. When a region contained fewer than 30 positive records, we nevertheless sampled 30 negative cases to ensure the minimal required representation of the randomly selected class. Using predicted probabilities and the true (hold-out) positive-negative labels, we calculated the area under the ROC curve (AUC) using the *pROC* package (Robin et al., 2021).

To account for stochastic variation from model fitting (e.g., randomness used by random forests in the tree-building process) and from the random sampling of negative records, we repeated the full workflow, including model fitting, variable importance estimation, LOOCV, and AUC calculation, across 10 random number seeds (values 0 to 9). For each region, we then calculated the mean AUC across the 10 random seeds, as well as the mean relative importance of each predictor.

To assess whether model performance differed among life forms (i.e., amphibians, birds, mammals, reptiles, and fish), we also calculated life-form-specific mean AUC values by subsetting predictions of the species from each group. Significant differences in performance across groups were assessed using a Kruskal-Wallis test (McKight and Najab, 2010).

### Exploratory analyses

We complemented the performance assessment with exploratory analyses that examined (i) the relative contribution of species traits versus spread history, (ii) whether spread-history predictor importance declines with geographic distance overall, and (iii) whether the strength of this distance-importance relationship, and overall model performance, varies with the global connectedness of focal regions.

First, we compared the performance of models using trait-only predictors with models using spread-history-only predictors. This comparison helps determine whether arrival in a focal region is better explained by intrinsic taxon characteristics (e.g., life form, habitat, diet, body size, and native-range attributes) or by prior occurrence elsewhere (years-since-arrival predictors). For each focal region, we fitted two additional random forest models: one including only trait predictors (life form, habitat, diet, body size, and native-range characteristics) and one including only years-since-arrival predictors. Each model was repeated across 10 random seeds, and mean AUC values were calculated per region for comparison.

Second, we tested for an overall geographic signal in the importance of spread-history predictors. Given the geographically structured nature of non-native species spread networks (Capinha et al., 2023), we expected spread-history predictors referring to regions closer to the focal region to be more informative. For each focal region–predictor-region combination, we calculated the shortest great-circle distance between regions (km; R Core Team, 2024) and paired it with the mean importance of the corresponding spread-history predictor (averaged across model repetitions). We then quantified the overall association between predictor importance and inter-regional distance (log-transformed) using repeated-measures correlation (Bakdash and Marusich, 2017), treating the focal region as the grouping factor to account for multiple observations per region.

Third, we tested whether overall model performance was associated with the global connectedness of focal regions. We expected mean performance to be lower in more globally connected regions because introductions are likely to draw from a broader, less geographically constrained pool of sources (Dawson et al., 2017; Hulme, 2021). We used total import volume in 2015 (i.e., the target reference) or, when unavailable, the closest available year as a proxy for connectedness, extracted from the Correlates of War Trade dataset (Barbieri et al., 2009). We then tested for an association between mean regional AUC and import volume using beta regression, after rescaling mean AUC to 0-1 to meet distributional requirements (Cribari-Neto and Zeileis, 2010).

### Predictions of species future arrival

We predicted the probability of future arrival for all species in each focal region using the ten replicate models fitted for that region and predictor values representing “present-condition” states. Specifically, we held species-trait predictors constant and updated spread-history predictors through 2015. We then generated predictions with each replicate model and calculated the average predicted probability across replicates. To restrict inference to regions achieving acceptable predictive performance, we retained only focal regions with a mean AUC ≥ 0.75.

For each of the two models with distinct retrospective calibration windows, we identified the species with the highest mean predicted probabilities of future arrival in each focal region. To ensure relevance for prevention planning and prioritisation, we retained only species for which we found no evidence of introduction in the focal region up to November 2025, based on a review of the literature and biodiversity-observation datasets. From this filtered set, we report the top 10-ranked species per region. We chose this cutoff (n = 10) for practical reasons, reflecting the resources available for verifying current introduction status.

For regions covered by both forecasting horizons (i.e., 20-year and 30-year models), we pooled the top-ranked species from both because we had no objective basis for prioritising one horizon over the other. For regions covered by a single horizon, we analysed the corresponding top-ranked list. We then examined species-frequency distributions across continents (ESRI, 2024) and taxonomic groups, quantified pairwise compositional similarity among regional lists using Simpson similarity (Koleff et al., 2003), and visualised broad inter-regional structure using multidimensional scaling (MDS) (Gower, 1966).

## Results

### Data summary

After excluding marine species and species with missing trait data, a total of 1,931 species were kept, of which 100 were amphibians, 936 birds, 446 fishes, 247 mammals, and 202 reptiles.

### Model performances

Models based on the 30-year period achieved very high predictive accuracy, with 58 regions out of the 110 modelled (∼53%) reaching a 10-replicate model average AUC of 0.9 or higher, 92 regions over 0.8 (∼84%), and only 8 regions falling below the minimum defined threshold of 0.75 (∼7%; Fig. 2a). Variability of AUC values across replicates was low (max. SD = 0.06). Across species groups, performance was consistently high (Fig. 2b), with median AUCs across regions above 0.9 for all groups except fishes (median = 0.88). A Kruskal-Wallis test found no significant differences in AUC values among taxonomic groups (*p* = 0.09).

Models using a 20-year calibration window showed similar performances. A total of 43 regions (∼56%) reached an average AUC of 0.9 or higher, 67 regions were above 0.8 (∼87%), and 8 regions were below the 0.75 threshold (∼10%; Fig. 2c). Variability among replicate models again remained low (max SD = 0.07). Across species groups, median AUCs were above 0.9 for amphibians, birds, and mammals, and above 0.85 for fishes and reptiles (Fig. 2d). We found no significant AUC differences between taxonomic groups (Kruskal-Wallis, *p* = 0.22).

Models jointly using predictors representing species traits and spread history substantially outperformed those based on either subset alone, for both 30-year and 20-year calibration periods (Fig. 2e,f; Kruskal-Wallis, *p* < 0.01). Between the individual subsets, models using spread history alone achieved significantly higher AUC values than those based only on species traits (Kruskal-Wallis, *p* < 0.01).

### Drivers of predictor importance and model performance

Repeated-measures correlation revealed a statistically significant but weak negative relationship between the relative importance of predictor regions (representing species spread history) and their geographical distance to the focal country (30-year: *r* = -0.12, *p* < 0.001; 20-year: *r* = -0.09, *p* < 0.001), suggesting that nearby regions are generally, but weakly so, more informative predictors of species’ future arrival.

In addition, we found a significant negative association between model performances (i.e., mean AUC value) and trade import volumes of the focal regions (30-year response: *p* = 0.044; 20-year response: *p* = 0.046), indicating that regions with higher global trade connectivity are more difficult to predict.

### Predictions of species future arrival

Correlation between predicted probabilities of regional arrival from the 30-year and 20-year models was very high (mean *r* across regions = 0.88). Across regions, the top 10 species (pooled across both forecasting horizons and restricted to models with mean AUC ≥ 0.75) were predominantly birds (n = 259; 59.4%), followed by fishes (n = 110; 25.2%), mammals (n = 27; 6.2%), reptiles (n = 26; 6.0%), and amphibians (n = 14; 3.2%). Taxa that ranked among the most likely to arrive in more than 15 regions included the birds *Phasianus colchicus* (n = 38), *Acridotheres tristis* (n = 30), *Amandava amandava* (n = 29), *Colinus virginianus* (n = 26), *Corvus splendens* (n = 19), *Lonchura malacca* (n = 19), *Alectoris chukar* (n = 17), *Psittacula krameri* (n = 16), *Lonchura oryzivora* (n = 15), *Numida meleagris* (n = 15), and *Pycnonotus jocosus* (n = 15), as well as the mammal European rabbit *Oryctolagus cuniculus* (n = 19). Several species were also predicted to arrive in regions spanning five to six continents, including birds *Amandava amandava* (n=6), and *Corvus splendens*, *Crithagra mozambica, Lonchura oryzivora*, *L. punctulata*, *Numida meleagris*, *Psittacula eupatria, P. krameri*, *Pycnonotus cafer*, and *P. jocosus* (all n= 5), fishes *Coregonus peled* and *Oreochromis mossambicus* (both n = 5), and a mammal (*Oryctolagus cuniculus;* n = 5).

At the taxonomic group-level, birds and fishes showed the broadest predicted arrival risk, with multiple species projected to arrive across many regions and across multiple continents, particularly in Asia, Central and South America and Europe (Fig. 3). In contrast, amphibians and reptiles showed patchier patterns that were more often restricted to single continents. Mammals showed intermediate patterns: a subset of species (e.g., *Oryctolagus cuniculus*, *Dama dama*) were predicted across several continents, whereas others remained geographically concentrated. Overall, for most species, predicted hotspots were concentrated within a few continents rather than being globally uniformly distributed, indicating strong regional structuring in predicted arrival risk across taxonomic groups.

**Figure 3.**
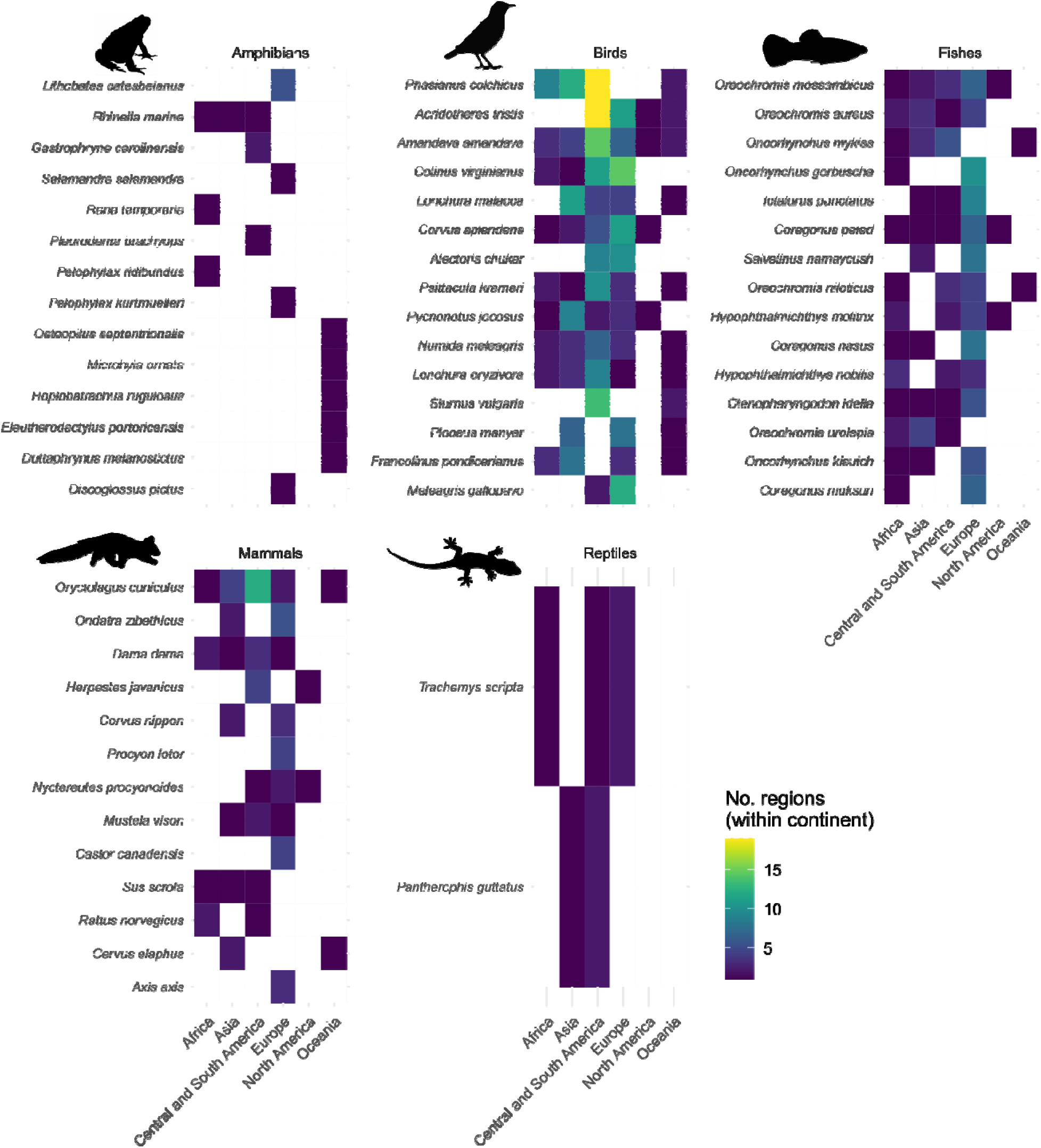
Species with the highest predicted spread potential by taxonomic group. Heatmaps show, for each taxonomic group, the species with the highest-ranked predicted probabilities of arrival across the greatest number of regions, pooling predictions from models trained on both retrospective windows (the 30-yr period, 1986–2015, and the 20-year period, 1996–2015). Within each taxonomic group, species were ranked by the total number of distinct regions in which they were predicted to arrive in the near future. Up to 15 species per taxonomic group are shown; ties at the cutoff were omitted, so some panels include fewer than 15 species. Columns represent continents, rows represent species, and tile colour indicates the number of regions within each continent where the species was predicted with the highest probability of introduction. White tiles indicate zero regions. In total, 102 regions are shown, representing those that met the minimum model performance threshold (AUC ≥ 0.75) for at least one retrospective window.

The MDS ordination of compositional similarity among high-risk regional species pools (top 10 predicted future arrivals per region) (Fig. 4) indicates moderate continental structuring with a pronounced zone of cross-continental overlap. Central and South American regions form a cohesive and distinct group. European regions also show a tendency to cluster, although several extend into the central overlap zone. Asian regions are comparatively dispersed, spanning the centre, with weak separation between Middle Eastern regions and East/Southeast Asian regions. North America (including the United States, Canada and Mexico) does not form a discrete cluster and instead falls mainly within the central overlap, intersecting with both European and Asian regions. Oceania is likewise widespread, with some Pacific islands (e.g., Guam and the Northern Mariana Islands) plotting nearer to the Central/South American cluster, while Australia occurs closer to Middle Eastern regions. Several regions (e.g., Hong Kong, Japan, the United States, Hawaii, China, Mauritius and Germany) plot near the centre, consistent with more “cosmopolitan” high-risk assemblages.

**Figure 4.**
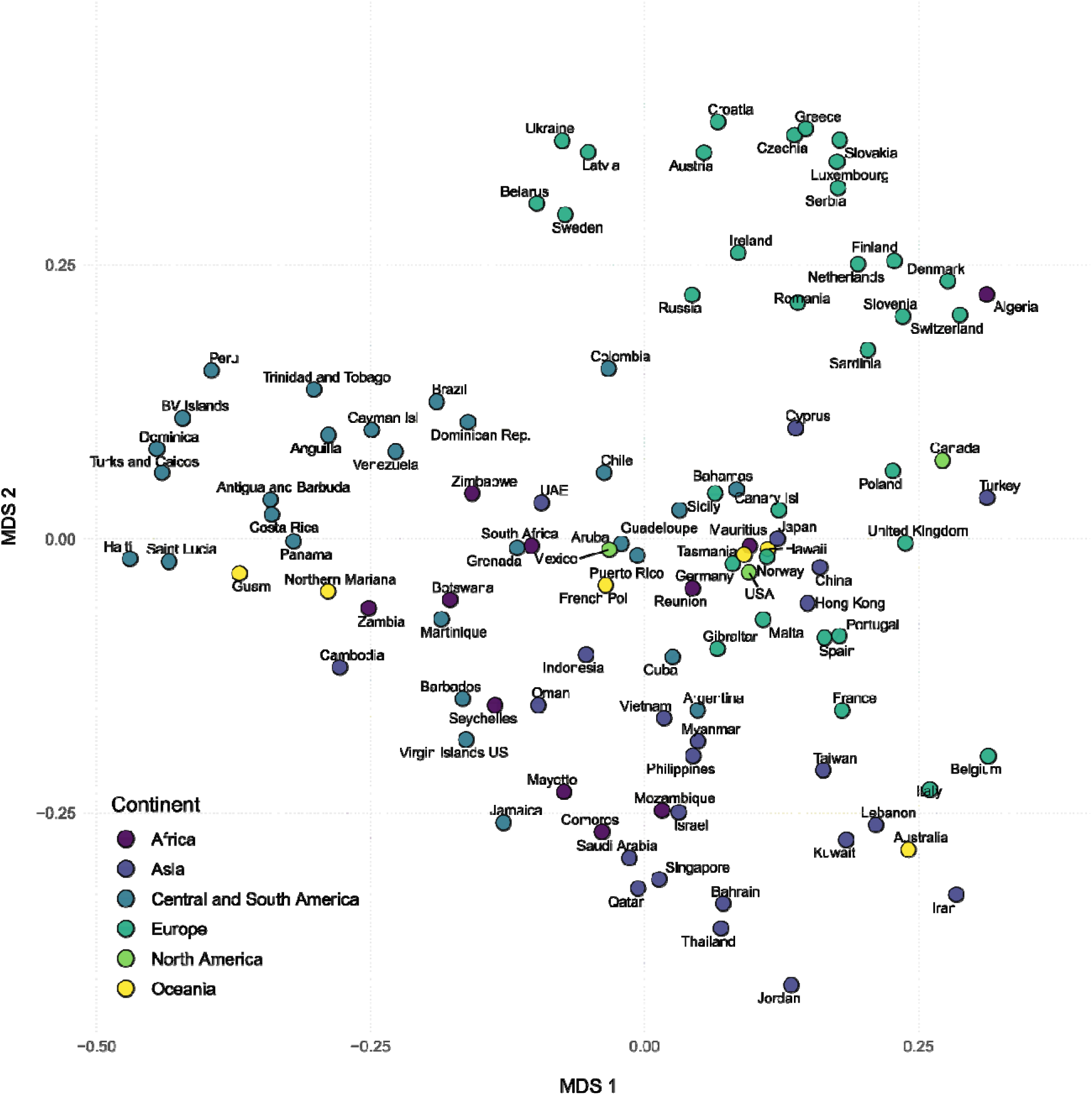
Multidimensional scaling (MDS) ordination of compositional similarity among regional high-risk assemblages (top 10 predicted future arrivals per region). Points represent 102 regions, corresponding to the subset of 110 modelled regions that achieved AUC values above the minimum performance threshold; colours indicate continent.

## Discussion

Our region-specific models produced consistently high predictive performance for near-future first records of non-native terrestrial and freshwater vertebrates worldwide, with most focal regions achieving strong discrimination (AUC >0.9 for more than half) across both retrospective calibration windows. Predictive performance also remained high across taxonomic groups. Together, these results indicate that near future spread potential is sufficiently well captured by spread-history and trait information to support operationally useful, region-level watchlists for proactive surveillance and prevention.

A central implication of these results is that, for many regions, the ability to anticipate likely “next arrivals” is high when using information that broadly mirrors what biosecurity practitioners and expert panels typically consider: where a species has already invaded previously, and the factors that may influence transport, establishment and early spread. In this sense, our framework can be interpreted as a scalable, repeatable analogue of expert horizon scanning, produced region-by-region and for hundreds to thousands of candidate species at once. This is particularly relevant because capacity for expert elicitation and sustained horizon scanning is unevenly distributed across the globe (Early et al., 2016; IPBES, 2023), as the number of potential invaders continues to rise (Seebens et al., 2025). A practical strength of our approach is therefore its deployability for individual regions, each receiving an internally consistent ranking of likely arrivals that can be updated as new first records accumulate and are paired with local feasibility/impact assessments to guide prioritisation.

Across regions, we identified a set of taxa repeatedly ranked among the most likely near-future arrivals. Many of the most recurrent taxa are already widely introduced and/or invasive, including frequently traded or transported birds such as the common pheasant (*Phasianus colchicus*), common myna (*Acridotheres tristis*) or the red avadavat (*Amandava amandava*) (Dyer et al., 2017), as well as freshwater fishes linked to aquaculture and stocking (e.g., Mozambique tilapia (*Oreochromis mossambicus*) and peled whitefish (*Coregonus peled*) (Froese and Pauly, 2025), and widely moved mammals such as the European rabbit (*Oryctolagus cuniculus*) (Long, 2003). The recurrence of these species across many regions suggests the models are capturing a realistic propagule-pressure gradient driven by regions where species have already become established and serve as hubs for further human-mediated redistribution. This interpretation is reinforced by model comparisons indicating that spread history provides a stronger signal than traits when considered alone: once a species has spread broadly, it carries a strong imprint of repeated introduction opportunity and potential for secondary dispersal. From a management perspective, species that repeatedly rank highly across regions are plausible candidates for coordinated, multi-region prevention and early detection.

At the same time, the predicted high-risk lists showed pronounced regionalisation, with some focal regions characterised by comparatively cosmopolitan sets of likely arrivals and others showing stronger continental clustering. This pattern is consistent with the view that near-term invasion risk reflects the interaction between global transport opportunity and regional-scale filters, including environmental matching and the composition and intensity of dominant introduction pathways (Lombaert et al., 2010; Bertelsmeier et al., 2018; Capinha et al., 2023). The compositional overlap observed among subsets of regional watchlists therefore indicates scope for coordinated, multi-jurisdiction prioritisation (e.g., joint horizon scanning exercises or shared candidate pools), particularly where cross-border connectivity and pathway structure are strongly shared. The EU Regulation 1143/2014 on Invasive Alien Species and its periodically updated Union list of priority species (EU, 2014) provide one illustrative example of such an approach, although similar rationales apply wherever invasion risks are demonstrably shared across borders.

Comparisons among different models showed that spread-history information was a stronger predictor than species traits when considered alone, but that the best performance was obtained when both were combined. This hierarchy is intuitive for short-horizon forecasting: where a species has already established often captures propagule pressure, pathway access, and “bridgehead” dynamics more directly than intrinsic traits (Bertelsmeier et al., 2018; Capinha et al., 2023). Traits nevertheless still add information, likely because they act as proxies for exposure to human transport pathways (e.g., association with trade, commensalism) and establishment and early spread capacity once introduced (e.g., generalism) (Blackburn et al., 2011; Allen et al., 2017; Gippet and Bertelsmeier, 2021). The superior performance of the combined (spread-history–trait) model therefore indicates that invasion risk is best predicted by an interaction between opportunity (history/connectivity) and capability (traits), and that neither information stream alone adequately represents the full invasion process.

A notable result is that model performance declined with increasing import volumes of focal regions, indicating that highly connected regions are more difficult to forecast. This pattern is consistent with evidence that international trade increases colonization pressure and is a major driver of biological invasions by expanding the pool of potential source regions and increasing the diversity of introduction pathways (Hulme, 2021). In such regions, arrivals may be drawn from many sources (including distant hubs) and via multiple pathways, so the risk signal contained in any single source–recipient relationship is diluted (Capinha et al., 2023; Montgomery et al., 2023). Consequently, identifying the most likely future arrivals poses a greater challenge in highly trade-connected regions and may require more explicit representation of connectivity and pathways to improve discrimination. This suggests that expert-based horizon scanning and risk assessments in highly trade-connected regions will likely face comparable challenges, because introductions may arise from a broader and more heterogeneous set of sources.

Beyond differences among regions in overall forecasting, we also evaluated how source-region informativeness is structured geographically and found a significant tendency for spread-history predictors from geographically closer regions to be more informative than those from distant regions. This is consistent with the idea that, once species enter a continent or biogeographic realm, subsequent spread and detection often cluster among neighbouring regions due to secondary spread and regional transport. However, the magnitude of the relationship was small, suggesting that geographic proximity captures only part of the connectivity that shapes inter-regional exchange. One interpretation is a two-stage process in which occasional long-distance introductions place species into new areas, followed by further spread that is more regionally clustered. Under this structure, proximity remains informative on average, but a smaller set of influential source regions, sometimes non-adjacent, can still contribute disproportionately. Moreover, neighbouring regions are unlikely to be uniformly informative, as their influence should vary with transport intensity, exchange structure, and the pool of present non-native species. This motivates the identification of “effective neighbours”, i.e., source regions that matter because of functional connectivity rather than adjacency. While recent studies have identified candidate drivers of this connectivity (e.g., commodity-specific trade flows; Oliveira et al., 2023; Zhang et al., 2024), such inputs and system-level knowledge are not consistently available at a global scale for multispecies forecasting. Our approach, therefore, uses invasion history as a scalable proxy for connectivity, with pathway- or trade-informed refinements where data permit.

### Limitations and future directions

Our approach carries limitations that should temper interpretation and guide future work. An important extension of this framework is to incorporate introduction pathways explicitly. Pathways can add mechanistic insight and may help explain why some non-adjacent regions function as influential “effective neighbours”, as well as why highly connected regions are harder to predict. However, pathway data remain sparse and uneven across taxa and regions, especially at the global scale and for large multispecies sets. Existing syntheses and linked-database approaches (e.g., Saul et al., 2017) provide valuable starting points but currently cover only a fraction of the species for which proactive forecasting would be useful. Therefore, a realistic near-term strategy may be to treat pathway information as an “add-on layer” where data exist (e.g., for priority taxa or well-characterised pathways), while continuing to rely on spread history and traits as a scalable baseline for broad forecasting.

Our approach also implicitly assumes that the near future resembles the recent past in terms of pathways, trade structure, regulations, and reporting. This assumption can be violated by major policy changes, rapid market shifts, conflict, infrastructure development, or new biosecurity measures, any of which can restructure introduction routes and invalidate extrapolation from historical patterns. For example, Reino et al. (2017) showed that the EU’s 2005 wild-bird import ban abruptly rewired global bird-trade routes, rather than simply reducing it everywhere. In addition, first-record data can be an imperfect proxy for true arrival because of gaps, detection and reporting delays and variation among regions in recording effort. These issues are particularly acute for the most recent years, where reporting lags can create apparent slowdowns and reduce the reliability of very-near-present calibration windows. Although our approach partially addresses this by ending calibration in 2015, residual bias and regional heterogeneity in detectability likely remain.

Further, our models do not consider emerging non-native species, i.e., non-native species that have not previously been recorded elsewhere as non-native. A substantial fraction of recent non-native species records globally involves such emerging species (Seebens et al., 2018), implying that history-based watchlists will systematically miss certain parts of future arrivals. This limitation underscores the need to pair quantitative forecasts with complementary approaches that can detect novelty. Predicted probabilities and rankings should therefore be interpreted as relative arrival likelihoods within the candidate pool represented in the underlying databases and trait coverage. Accordingly, they are best used to prioritise among known potential invaders rather than as exhaustive predictions of all future invaders.

## Conclusions

Our results show that data-driven, near-term regional forecasting of non-native vertebrate spread can be operationally useful across broad spatial and taxonomic extents. Performance was generally high, supporting the development of regularly updated, region-specific watchlists to inform surveillance and prevention. Notably, many widely introduced and well-known invaders still emerged among the most likely near-future arrivals across regions, indicating substantial unrealised spread potential and continued scope for prevention and early detection. At the same time, reduced performance in highly trade-connected regions and the limited explanatory power of geographic proximity for identifying informative source regions suggest that forecasting future arrivals may remain challenging in some contexts. Integrating pathway data where available and accounting for detection lags should further strengthen the robustness and interpretability of regional arrival forecasts.

## Acknowledgments

This work acknowledges funding from Fundação para a Ciência e Tecnologia (FCT) through InvaSTOP grant (https://doi.org/10.54499/2023.12533.PEX) and through funds to CEG/IGOT Research Unit (UID/00295/2025; https://doi.org/10.54499/UID/00295/2025).

## Data Availability Statement

Data and code will be publicly available upon peer-reviewed publication.

