## Appendix 1 - Supplementary Material for "Data-driven forecasts of regional arrivals of non-native vertebrates worldwide"

### Supplementary materials for: Data-driven forecasts of regional arrivals of non-native vertebrates worldwide

#### Supplementary figures

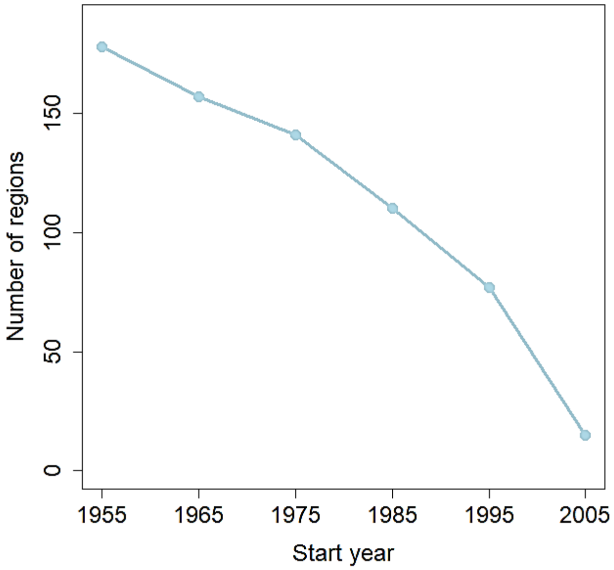


**Figure S1.** Relationship between the start year of the retrospective window (ending in 2015) and the number of regions with ≥5 first records (minimum for model calibration), evaluated in 10-year cumulative increments, illustrating the trade-off between recency and model coverage.

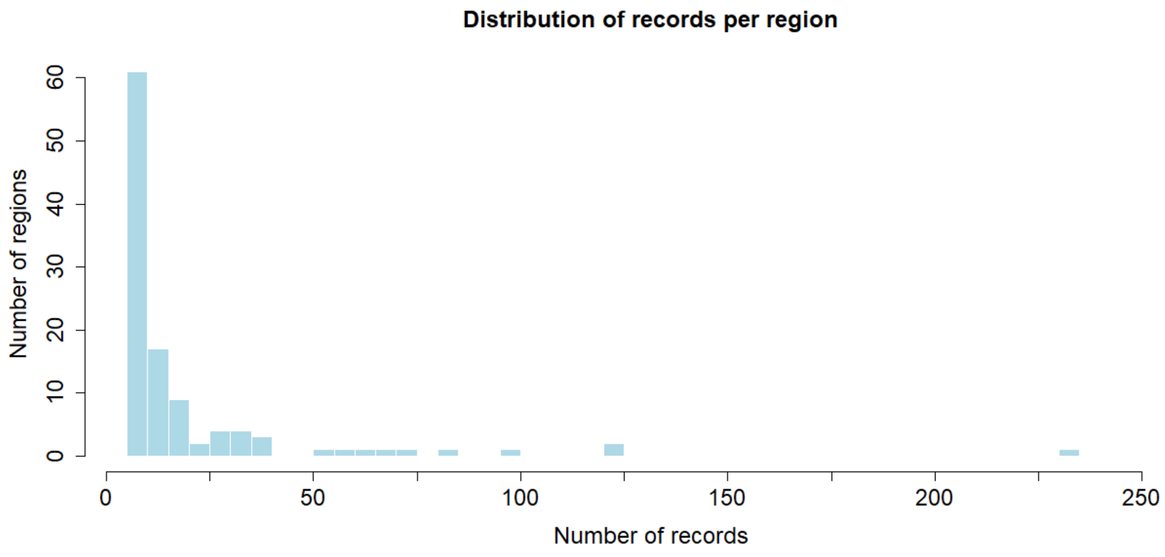


**Figure S2.** Histogram of first-record counts across regions for the 30-year (1986–2015), showing regions meeting the minimum threshold of ≥5 first records for model calibration.

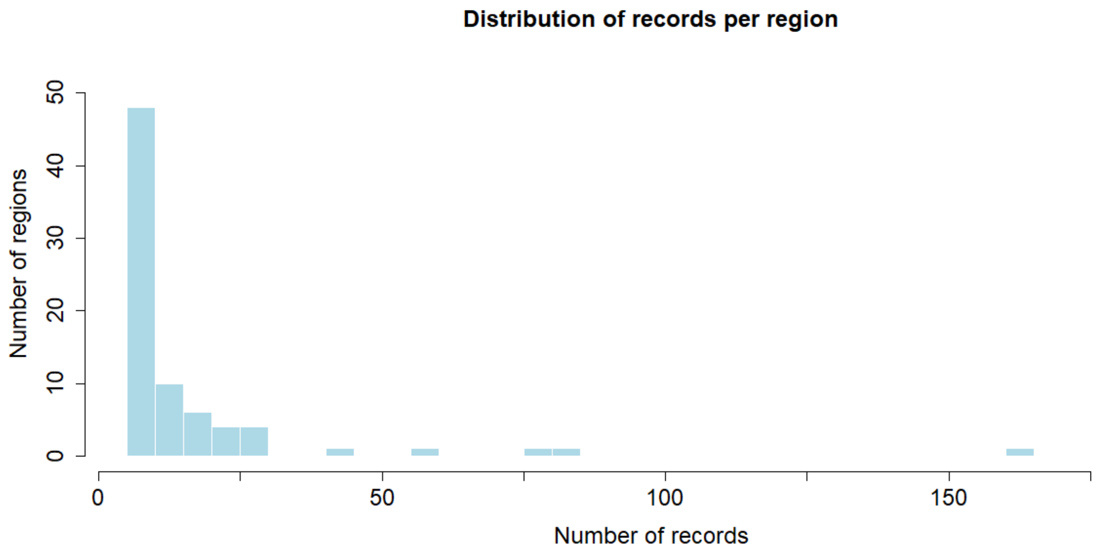


**Figure S3.** Histogram of first-record counts across regions for the 20-year (1996–2015), showing regions meeting the minimum threshold of ≥5 first records for model calibration.
